# Cortical cooling reveals a role for visual cortex in generating visual responses in auditory cortex

**DOI:** 10.64898/2026.06.10.731095

**Authors:** Rebecca H.C. Norris, Stephen M. Town, Katherine C. Wood, Jennifer K. Bizley

## Abstract

Multisensory integration is a fundamental feature of cortical processing, yet the functional pathways that deliver visual signals to the auditory cortex remain poorly understood. While anatomical studies reveal multiple candidate projection routes, demonstrating their causal contribution requires targeted manipulation of neural activity. Here, we used cortical cooling to reversibly inactivate the posteromedial lateral suprasylvian cortex (PMLS) and the adjacent area 21 to determine the functional role of higher-order visual areas in generating visual responses within the auditory cortex of the ferret.

Units responsive to sound, light, or combined audiovisual stimuli were found across all sampled auditory fields and cortical depths, with visual responses most prominent within the infragranular layers and the non-tonotopic secondary auditory cortex of the Anterior Ectosylvian Gyrus (AEG). Cortical cooling induced robust, bi-directional, and stimulus-specific modulations of firing rates in AC. Approximately 50% of visually responsive units exhibited a significant decrease or complete elimination of visual activity during cooling, confirming a functional role for visual input from PLMS/area 21 to AC.

Surprisingly, cooling also revealed circuit-level complexities: a subset (∼5%) of units showed enhanced or newly emergent visual responses during inactivation, suggesting that PMLS/area 21 normally exerts a gating influence over alternative visual pathways. Furthermore, contrary to feedforward anatomical predictions, neurons in the AEG—the region most heavily innervated by the cooled visual areas —were less frequently impacted by cooling than those in PEG. Together, these findings demonstrate that higher visual areas causally shape cross-modal processing in the auditory cortex through a complex mixture of direct excitation and network-level modulation.

## Introduction

Multisensory integration is a fundamental property of mammalian sensory systems. Focusing on the integration of visual signals into auditory processing, effects emerge within the early sensory cortices, across species including mice (Morrill & Hasenstaub, 2018), rats (Wallace et al., 2004), ferrets (Bizley et al., 2007) cats (Meredith & Allman, 2009), macaques (Ghazanfar et al., 2005; Kayser et al., 2008) and humans (Besle et al., 2008).

Anatomical studies provide a plethora of potential sources of this visual input to auditory cortex (AC) including feedforward inputs from subcortical structures, lateral or feedback connectivity from visual cortex, and feedback connections from higher cortical areas such as the parietal cortex (Bizley et al., 2007; Bizley & King, 2009; Budinger et al., 2006, 2008; Lee & Winer, 2011). However, despite an abundance of possibilities it is unclear which of these pathways subserve the visual responses observed electrophysiologically. For example, despite a robust anatomical projections from auditory cortex to area 21 in the ferret, one study found minimal evidence for suprathreshold audiovisual integration (Allman et al., 2008).

Demonstrating a functional role for an anatomical projection therefore requires targeted manipulation of neural activity. Very recent studies have used this approach to demonstrate a functional role for visual cortex in visual responses in auditory cortex; optogenetically silencing V1 in the mouse suppressed visual evoked firing in AC (Olsen & Hasenstaub, 2025) and cooling visual cortex in the ferret eliminated the impact of visual stimuli on the LFP in auditory cortex (Atilgan et al., 2018).

While in the mouse V1 directly innervates A1, in the ferret (as in the primate) direct connections from V1 are very sparse, while connections from secondary visual areas to secondary auditory cortex are more prominent. In the ferret, connections between secondary visual cortex and auditory cortex occur in an ordered and field specific manner; Area 20, a visual form area, provides strong innervation to the Posterior Ectosylvian Gyrus (PEG), where tonotopic secondary auditory cortex is located, whereas area PMLS and area 21 strongly innervate the Anterior Ectosylvian Gyrus (AEG) where non-tonotopic secondary auditory cortex is located.

In this study we took advantage of the known cortical anatomy of the ferret and its multiple well-spread auditory and visual cortical fields to assess the impact of silencing one of the visual cortical sources of input to auditory cortex. We focused specifically on PMLS/area 21 which is known in the ferret to project directly to non-tonotopic secondary AC (AEG). PMLS is considered broadly analogous to primate area MT (Lempel & Nielsen, 2019; Manger et al., 2008; Payne, 1993) (note PMLS has been referred to in the ferret visual cortical literature as both the Suprasylvian Visual area (SSY) and the Posterior Suprasylvian field, PSS). To manipulate cortical activity we used cortical cooling of the gyral surface of PMLS and the adjacent field 21. Cooling allowed us to reversibly manipulate visual cortex while simultaneously recording neural activity in auditory cortex.

The advantage of cortical cooling over alternative methods such as physical lesions or pharmacological inactivation, is that cortical cooling is acute and reversible on a fairly rapid timescale, limiting the potential for compensation by other brain regions and allowing within-subject comparison of alternating control and inactivation time periods. Additionally, cortical cooling allows the manipulation of large areas of tissue that can be difficult when using genetic tools or optical methods (Lomber et al., 1999; Wood et al., 2017). Previous studies have used cortical cooling to investigate sensory and decision-making neural processing (Antunes & Malmierca, 2011; Nakamoto et al., 2010; Orton et al., 2012).

We tested the effects of reversible inactivation of PMLS/area 21 on activity in ferret AC while anesthetised animals were presented with a series of auditory, visual or combined audio-visual stimuli. Performing these recordings while animals were under anaesthesia allowed us to eliminate potential effects of attention and motor actions on multisensory processing (Bimbard et al., 2023; Olsen & Hasenstaub, 2025; Oude Lohuis et al., 2024). We expected to reproduce previous findings of visual and audio-visual responses in ferret auditory cortex, and we predicted that at least some portion of these responses would be eliminated or reduced by cooling PMLS/area 21. Consistent with our predictions, we found visual and audiovisual (AV) responses across all fields of ferret AC. While many of these responses were indeed reduced by cooling PMLS/area 21, we additionally found units where visual or AV activity was unimpacted by cooling, suggesting that other pathways also influence AV responses in AC. We also found a small percentage of units where visual responses emerged during cooling, suggesting that under normal conditions PMLS/area 21 inhibits some of these responses.

## Methods

### Animals

Subjects were six adult pigmented ferrets (*Mustela putorius*, female; 1–4 years) who underwent electrophysiological recordings under non-recovery anesthesia while visual cortical activity was modulated with cooling. These animals were naïve to any auditory tasks. Prior to the experiment, all ferrets were housed in groups of two to eight, with free access to high-protein food pellets and water, with regular otoscopic examination to ensure that both ears of the animals were clean and healthy. All experimental procedures were approved by the local ethical review committee and were carried out under license from the UK Home Office, in accordance with the Animals (Scientific Procedures) Act 1986.

### Electrophysiological recording

Anaesthesia was induced by a single dose of a mixture of medetomidine (Domitor; 0.022 mg/ kg/h; Pfizer) and ketamine (Ketaset; 5 mg/kg/h; Fort Dodge Animal Health). The left radial vein was cannulated and anaesthesia was maintained throughout the experiment by a continuous infusion of medetomidine (0.022 mg/kg/hr) and ketamine (5 mg/kg/hr), with atropine sulphate to reduce bronchial secretions (0.06 mg/kg/hr, C-Vet veterinary products) and dexamethasone to reduce cerebral oedema (0.5 mg/kg/hr, Dexadreson, Intervet UK) in Hartmann’s solution, supplemented with 5% glucose. The ferret was intubated, placed on a ventilator (Harvard Model 683 small animal ventilator; Harvard Apparatus) and supplemented with oxygen. Body temperature (38 °C), end-tidal CO2, and the electrocardiogram were monitored throughout the experiment. Experiments typically lasted between 36 and 60 hours.

To access the cortex for recordings, the animal was placed in a stereotaxic frame and the temporal muscles on both sides were retracted to expose the dorsal and lateral parts of the skull. A metal bar was cemented and screwed into the right side of the skull, holding the head without further need of a stereotaxic frame. On the left side, the temporal muscle was largely removed, and a craniotomy performed to expose the suprasylvian and ectosylvian gyri where visual and auditory cortices are respectively located. The dura was removed and the cortex covered with 2% agar. The animal was then transferred to a small table in a soundproof chamber (Industrial Acoustics, Winchester, UK).

Recordings were made using silicon probe electrodes (Neuronexus Technologies, Ann Arbor, MI) with either a single shank (16 channel; 100 μm site spacing; placed in VC), or a double shank (32 channel; 100 um site spacing; placed in AC). Electrodes were positioned so that they entered the cortex approximately orthogonal to the surface of the suprasylvian gyrus or auditory cortex. Neural recordings were obtained using TDT System III hardware (RZ2 data acquisition system) with custom written software in Open Project (Tucker-Davis Technologies, Alachua, FL) and MATLAB (Mathworks, Natick, USA).

At all sites recordings were made prior to, during and after cooling. Across all recording sites the mean loop temperature was ± SD =7.14 C (± 1.02, minimum = 5.70 °C). The brain was allowed to recover until it returned to within 3 °C of the pre-cooling temperature (minimum 25 minutes, mean ± SD 54.5 min ± 16 min).

### Cortical cooling

To reversibly inactivate visual cortex (VC) we cooled the surface of the brain using a cooling loop. The cooling loop was a modified, miniaturised version of the cooling loop developed by (Lomber et al., 1999), as previously described in (Wood et al., 2017). The cooling loop was constructed from 23 gauge stainless steel tubing which was bent to form a loop shape that ran dorso-ventrally posterior to the suprasylvian sulcus. In contrast to loops used in auditory cortex, where the shape formed a circular loop, for silencing PMLS/area 21 which are narrow fields that run dorso-ventrally within and adjacent to the posterior sylvian sulcus, we used designed the loop so that only one edge of the loop ran along the cortical surface forming a dorso-ventrally oriented bar, rather than a circle. A micro-thermo- couple, made from twisting together 30 AWG gauge (0.254 mm) PFA insulated copper and constantan wire (Omega Engineering Limited, Manchester, UK), was soldered to the base of the loop and secured with an epoxy adhesive. The thermocouple wire was soldered to a miniature female thermocouple connector (RS components Ltd, UK) and again secured with an epoxy adhesive.

In each experiment the cooling loop was supplied with ethanol from a reservoir via an FMI QV drive pump (Fluid Metering, Inc., NY, USA) controlled by a variable speed controller (V300, Fluid Metering, Inc., NY, USA). Ethanol was carried in FEP and PTFE tubing (Adtech Polymer Engineering Ltd, UK) from the reservoir to pump (FEP: 1.1 mm x 2 m, inner diameter x length) and then from pump to cooling loop (FEP: 0.8 mm ID x 2 m, PTFE: 0.5 mm x 2 m). Where necessary, tubing was bridged using two-way connectors (Diba Fluid Intelligence, Cambridge, UK).

To cool the ethanol prior to arrival at the cooling loop, 1 m of PTFE tubing was coiled within a Dewar flask (Nalgene 4150–1000, NY, USA) containing a mix of ethanol (100% concentration) and dry ice (BOC, UK). To maintain ethanol temperature within the tubing system, the length of tubing between the Dewar flask and cooling loop was minimized and insulated with silicon tubing. Ethanol from the loop was recycled into the reservoir. Cooling loops were positioned on the cortex with a microdrive.

The efficacy and spread of cooling was confirmed at the end of the experiment using a hypodermic needle temperature probe (Omega, Stamford, USA) at varying distances from the cooling loop, in both visual and auditory cortex. At each site, a micromanipulator was used to position the temperature probe in order to sample the temperature in cortex, in 500 μm steps to a depth of 2.5 mm from the cortical surface (these data were presented in Fig.2A (Wood et al., 2017)

During cooling, cortical temperature in PMLS/area 21 (VC) was reduced to between 6.0 and 10 C. At these temperatures, visual responses were reversibly suppressed, while simultaneously maintaining auditory cortical temperature and neural responses (Figure 1C). An electrode placed in visual cortex confirmed that the effects of cooling were restricted to within 500 um of the loop.

**Figure 1.**
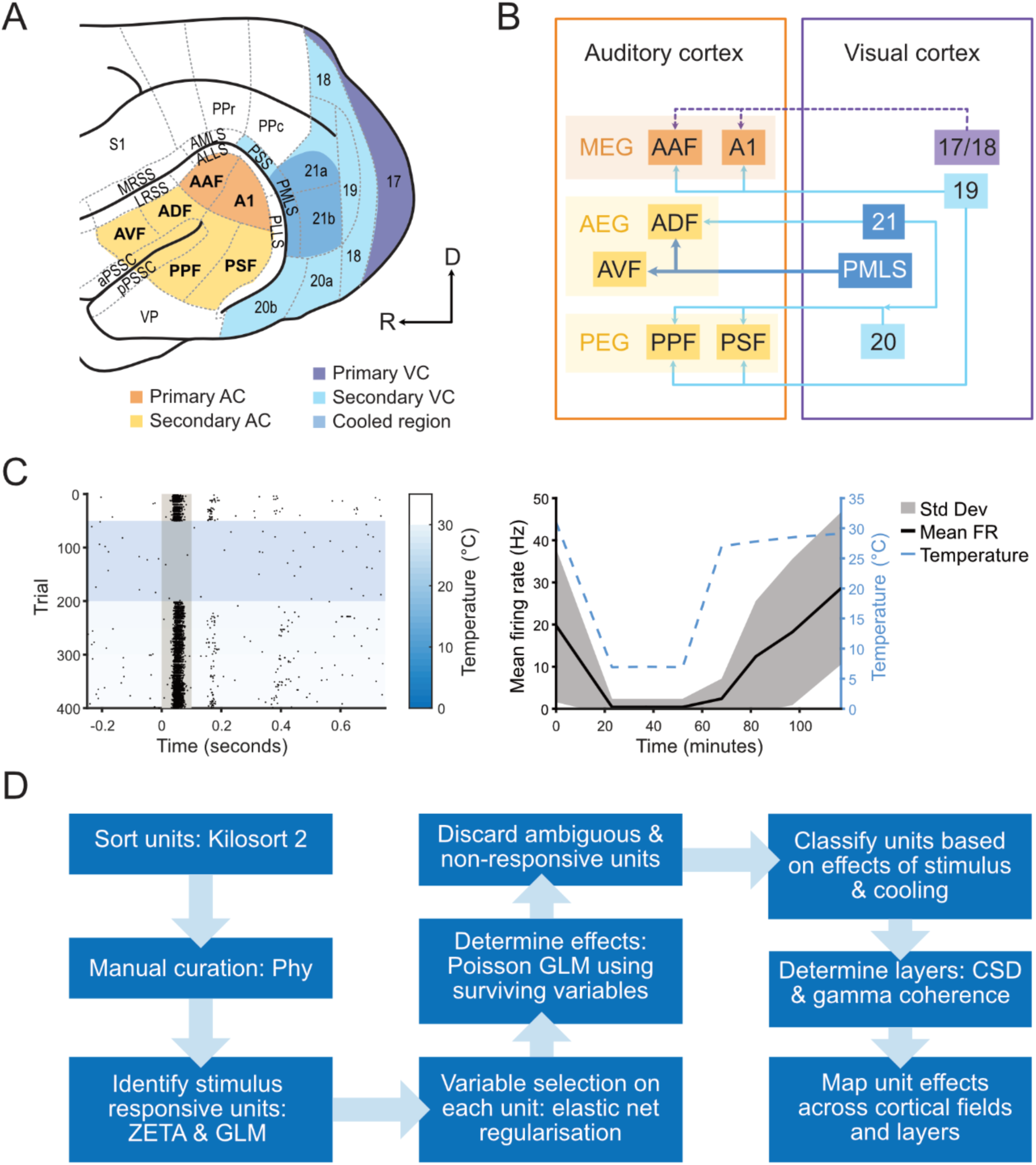
A) Diagram of the ferret auditory and visual cortices colour coded to show primary and secondary areas; the cooled region of extrastriate visual cortex covering PMLS and area 21 is marked in dark blue. A1, primary auditory field; AAF, anterior auditory field; ADF, anterior dorsal field; AVF, anterior ventral field; PPF, posterior pseudosylvian field; PSF, posterior suprasylvian field; VP, ventral posterior; a/pPSSC, anterior/posterior pseudosylvian sulcal field; M/LRSS, medial/lateral rostral suprasylvian sulcus; PSS, posterior suprasylvian sulcus; PMLS posteromedial lateral suprasylvian area; PLLS, posterolateral lateral suprasylvian area; S1, primary somatosensory; PPr/c, rostral/caudal posterior parietal; 17–21, visual areas 17–21. B) Schematic of known connections between ferret visual and auditory cortices, dotted lines indicate sparse connections, solid line colour depth indicates density of connections. High level divisions of auditory cortex are AEG, anterior ectosylvian gyrus; MEG, middle ectosylvian gyrus; PEG, posterior ectosylvian gyrus. C) Spike raster (left) and mean firing rate (right) alongside cortical temperature of an example unit recorded from the cooled region of visual cortex before, during and after cooling showing reversible inactivation during cooling and recovery of firing after the cortex was allowed to return to normal temperature. D) Schematic of the analysis pipeline.

After cooling the brain was allowed to recover until it returned to within 3 °C of the pre-cooling temperature (minimum 25 minutes, mean ± SD 54.5 min ± 16 min).

### Stimuli

Auditory stimuli were white noise bursts, of duration 100 ms, cosine ramped with 5-ms duration at the onset and offset and low pass filtered below 22 kHz (finite impulse response filter <22 kHz, 70 dB attenuation at 22.2 kHz). Noise bursts were generated afresh on each trial in MATLAB at a sampling frequency of 48848.125 Hz and presented from one speaker contralateral to the recording site.

Visual stimuli were white LED light flashes from an LED positioned 30 cm from the contralateral eye/ferret’s head. All stimuli were 100ms in duration. Trials were of 3 types: auditory, visual or both auditory and visual stimuli, with the order of trial types pseudorandomly generated. Typically 50 repetitions of each stimulus were presented within a block at a given site / temperature.

### Data analysis

Continuous voltage signals were recorded at 24414 Hz, and filtered into low pass and high pass streams for local field potential (LFP, 10-500 Hz) and spiking analysis (from 300-5000 Hz). For recording sites in AC, high pass streams were concatenated across contiguous blocks from pre-cooling through cooling to post-cooling, then spike sorted using Kilosort 2 (Pachitariu et al., 2016).. Units yielded by Kilosort 2 were inspected and manually curated in a specialised graphical user interface (https://github.com/cortex-lab/phy).

We assessed recovery of units by comparing stimulus evoked firing rates (FR) for pre-cooled and post-cooled recording blocks. Units which recovered to a less than 20% difference from their pre-cooled FR during the post-cooled recording blocks were deemed to have ‘good’ recovery, while units that recovered to a less than 50% difference from their pre-cooling FR were deemed to have ‘fair’ recovery. Units which failed to meet either recovery criterion were excluded from analyses, similar to previous cooling studies, e.g., (Antunes & Malmierca, 2011). We were unable to assess recovery for units with very low firing rates (< 1.3 hz) using this method due to the outsize impact on FR contributed by single spikes, so these units were excluded if both pre-cool and cooled blocks or both cooled and post-cooled blocks were below 1.3 hz. We additionally excluded units which exhibited consistent increases or decreases in FR of more than 20% across pre-cooled to cooled to post-cooled blocks, as we considered this pattern more likely to reflect electrode drift than a reversible impact of cooling visual cortex. After excluding units that did not meet criteria for inclusion, we identified 221 multiunits and 59 single units which are analysed together here.

To categorise sorted units according to their modality response and the effect of cooling, we employed a two-step feature selection and regression strategy to analyse spike counts pre and post stimulus using bin sizes of 100ms and 350ms to capture fast as well as slow latency responses. First, an Elastic Net regularized Poisson regression (alpha = 0.5, function lassoglm in MATLAB (The MathWorks Inc., 2022)) was applied to a predictor matrix containing a full factorial combination of auditory stimulation, visual stimulation, and cooling state, alongside a continuous covariate for trial number. Model hyperparameters were optimized using 10-fold cross-validation, and active variables were selected using the conservative one-standard-error (1SE) criterion. Second, the surviving predictors were entered into a final, unregularized Poisson Generalised Linear Model (GLM) (function fitglm in MATLAB). Using this feature-selection and re-estimation approach allowed us to ensure a sparse model architecture while preventing the parameter shrinkage bias associated with regularized coefficients.

To account for more complex patterns in firing rate, we additionally employed ZETA tests to classify sorted units as stimulus responsive. Our data showed a wide range of onset latencies for stimulus evoked firing, and the temporal pattern of spiking was sometimes impacted by cooling, making it less well suited to approaches that depend on an arbitrary bin size (e.g. t- tests, ANOVA, PSTH and our two-step approach described above). ZETA tests do not require binning, and the temporal resolution of a ZETA test is limited only by the spike density, which allows more accurate determination of instantaneous firing rate and peak response latencies (Montijn et al., 2021). Sorted units were classified as stimulus responsive if the result reached significance in either a GLME or a ZETA test. Onset latencies were based on the instantaneous firing rate (IFR) output of the ZETA tests, defined as the latency to half-peak height. To assess changes in stimulus-evoked firing rate during cooling compared to pre-cooling, we calculated the mean and standard deviation (SD) of each unit’s firing rate across the full pre-cooling block, in order to encompass each neuron’s dynamic range. This mean and SD was used to calculate z-scored firing rates to assess stimulus-evoked activity. Z-scored firing rates were compared between cooled and pre-cooled (control) blocks by subtracting the z-scored firing rate during cooled from the rate during control blocks.

### Localisation of recording sites

To determine which field recordings were made in we used information from photographs of the position of electrode penetrations and tonotopic maps constructed using the frequency response areas from auditory evoked units on each probe in response to pure tones, alongside the PSTHs of units in response to tones and noise bursts.

To determine which cortical layer our units were recorded in, we used the LFP streams to calculate and analyse local field potential (LFP) and current source density (CSD) profiles for each site. Data in the LFP streams were low-pass filtered below 150 Hz (zero phase shift, 4th order butterworth filter) and downsampled to 498 hz, following which 50Hz line noise was removed using a spectrum interpolation method implemented in the fieldtrip toolbox (Leske & Dalal, 2019; Oostenveld et al., 2011).We considered several factors to assign channels to supragranular (layers 1 - 3), granular (layer 4) and infragranular (layers 5 - 6) cortical layers. The key factors we used were: the pattern of CSD sources and sinks evoked in response to noise bursts and/or pure tones; the shape of the LFP traces in response to noise bursts and/or pure tones; the relative power in each of 3 bands of interest: theta (1-4Hz), alpha/beta (10-30 Hz), and gamma (50-150 Hz), and the pairwise channel by channel correlations in power in the gamma band; features were visualised using a custom graphical interface (https://github.com/LBHB/laminar_tools) as reported in (Hamersky et al., 2025).

## Results

We recorded the response of 280 neurons that met our criteria for inclusion, throughout primary and secondary auditory cortex to auditory and visual stimuli before, during and after cortical cooling of visual cortex (Figure 1). Replicating previous studies in the ferret and other animals, we revealed a diversity of response types, including auditory units uninfluenced by the visual stimulus through to units that responded only to the visual stimulus with a spectrum of responses in between.

Considering only units whose responses were sufficiently stable between pre- and post-cooling sessions (those that made a ‘fair’ or ‘good’ recovery, see methods for details) we asked whether silencing non-primary visual cortex (specifically areas PMLS and area 21) impacted auditory cortical activity.

Again, we saw a spectrum of effects, with some units’ activity unchanged by cooling (Fig.2A), while others were clearly modulated, showing elimination of the entire visual response (Fig.2B) or elements of it (Fig.2D), or even the emergence or strengthening of a visual response (Fig.2C,E). To classify the effects across all recorded units, we used a poisson GLM (with elastic net regularization) to determine, for each unit, its modality as either auditory (‘A’), visual (‘V’) or audiovisual (‘AV’, including auditory or visual responses with a cross-modal interaction as well as neurons that responded to both A and V) and to ask what the impact of cooling was.

**Figure 2.**
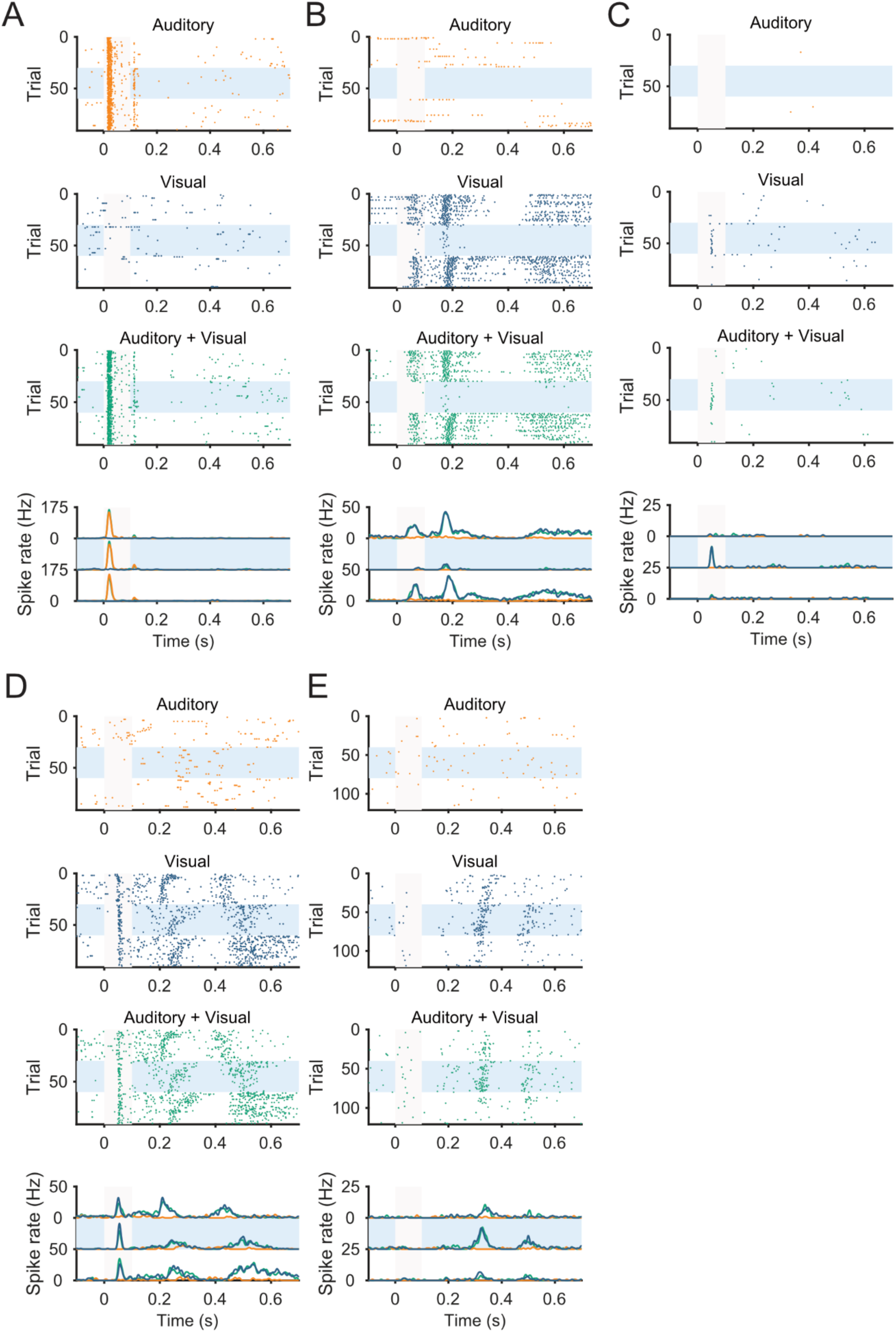
Example Data. Spike raster and peri-stimulus time histogram (PSTH) plots for five example units recorded in auditory cortex in response to auditory (yellow), visual (blue) and combined auditory and visual (green) stimuli. The stimulus duration is indicated by a grey bar. Trials are stacked from top to bottom in the order they were presented (stimulus types were presented in a pseudorandomised order during each time interval) pre, during (indicated by a blue box) and after cooling visual cortex. A) Example auditory multi-unit, showing no impact of cooling. B) Example visual single unit, showing decreased stimulus-driven firing during cooling. C) Example visual single unit, showing emergence of stimulus-driven activity during cooling. D) Example visual multi-unit showing a more complex impact of cooling: preserved onset activity and disrupted/delayed offset activity. E) Example visual unit showing increased stimulus-driven activity during cooling.

**Figure 3.**
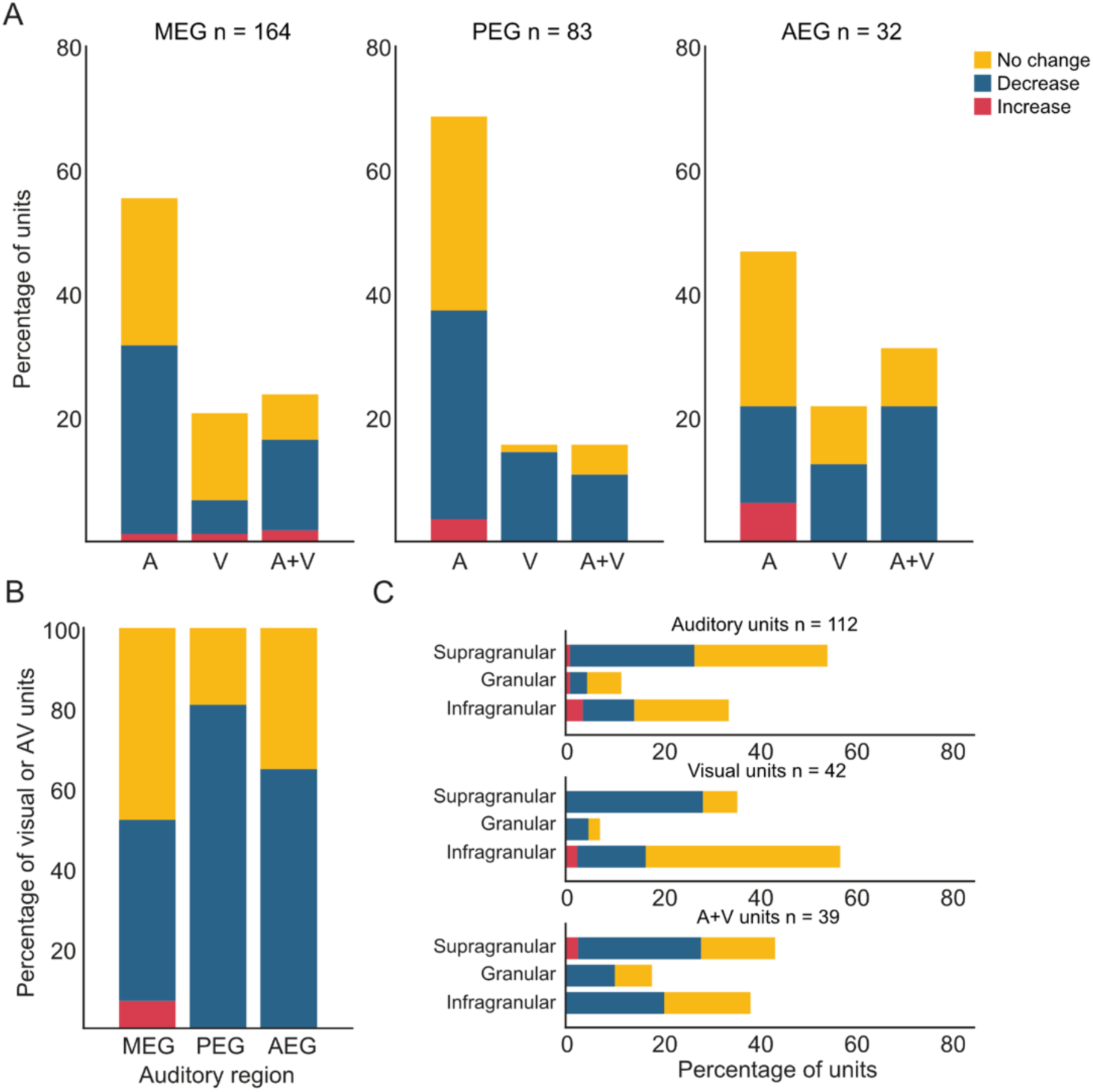
Summary of the effects of cooling. Bar plots of percentage of units coloured by impact of cooling: increased firing rate (red), decreased firing rate (blue), no change in firing rate (yellow). A) Percentages of units falling into three mutually exclusive stimulus responsivity categories: A, auditory only; V visual only; A+V, both auditory and visual, or any interaction effect. Data are divided by the recorded field into MEG, PEG and AEG. B) Summary percentages of only units showing visual responses or an interaction between auditory and visual responses across the same three main divisions of AC. Note that one unit is not included in this plot due to its field location being uncertain C) Summary percentages of units in each stimulus response category plotted by the layer the units were in. Note that the unit counts are lower for the plot due to being unable to accurately determine layer for some units included in plots A and B.

With this approach we identified 164 auditory, 54 visual and 62 auditory visual responses. We asked how these responses were distributed across the three subdivisions of auditory cortex; MEG (Middle Ectosylvian Gyrus); containing the primary fields A1 and AAF, PEG; containing secondary tonotopic fields PPF and PSF, and AEG; containing non-tonotopic secondary fields ADF and AVF. PEG had a significantly lower proportion of visual or multisensory units than AEG, while there was no significant difference in the proportions of visual/multisensory units between MEG and either PEG or AEG.

Across units, the bi-directional effects of cooling on firing rate were also captured, with 13 units (4.6%) increased, 148 units (52.9%) decreased, and 119 units (42.5%) uninfluenced by cooling. Figure 4A illustrates the impact of cooling on each category of unit, across the three major cortical subdivisions. Given the anatomical connectivity of PMLS and area 21, which most strongly connects to the AEG, we predicted that visual responses in this area would be most likely to be impacted by cooling. Contrary to this, visual neurons in all three areas were impacted by cooling, and those in AEG were significantly less likely to be impacted by cooling than those in MEG & PEG (although we note the relatively small sample size in AEG).

**Figure 4.**
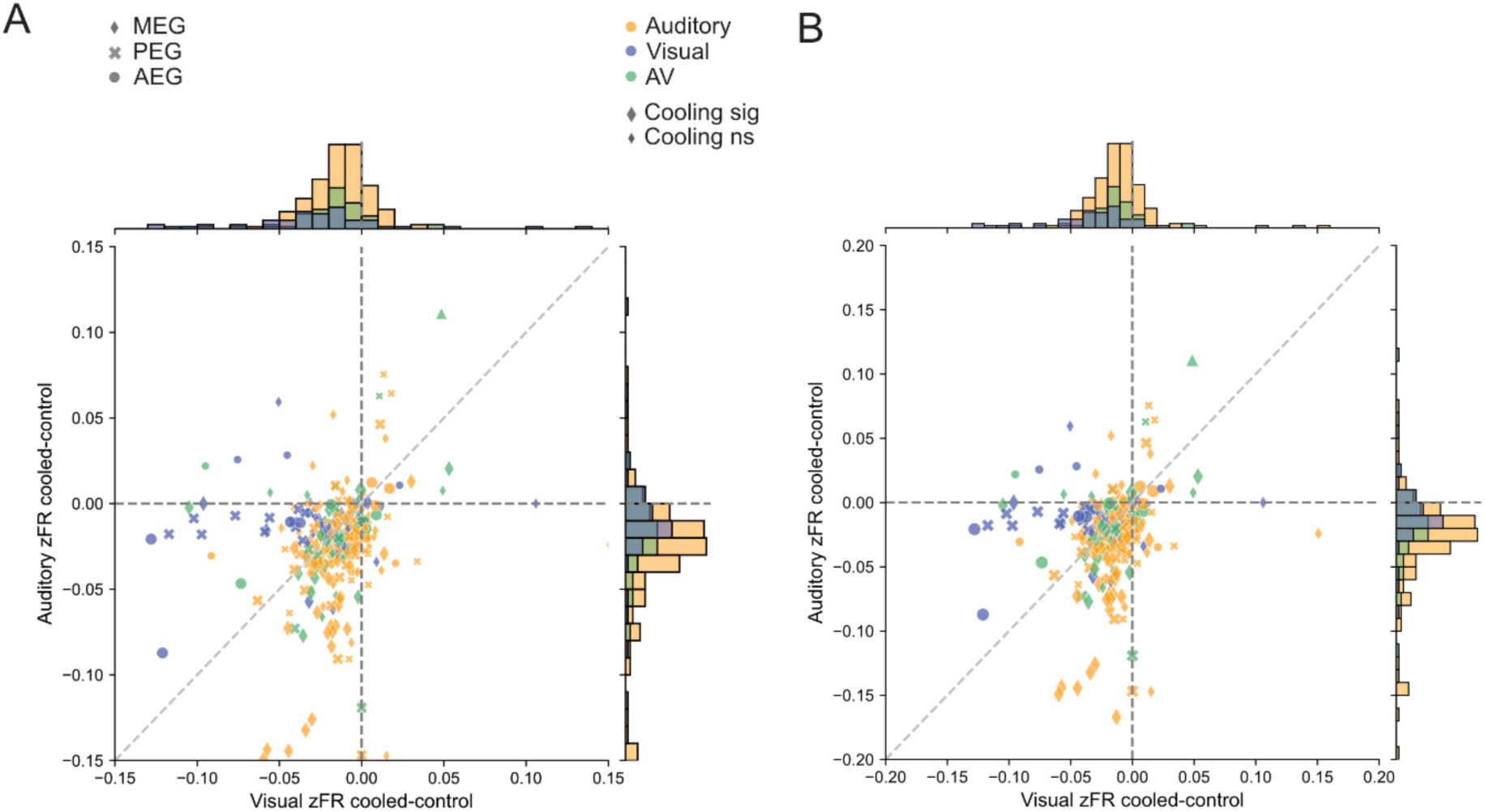
Changes in stimulus evoked firing rate from pre-cooled to cooled blocks. A) Scatter and marginals plots of the difference between z-scored firing rates for the period 0 - 350ms following stimulus onset in the control (pre-cooled) block compared to the cooled block. A) Scatter and marginal plots of the difference between control and cooled z-scored firing rates for auditory trials (y axis) against visual trials (x axis). Points are colour coded according to whether the unit was classified as auditory responsive (yellow), visual responsive (blue) or AV (green). Marker shape indicates the field that each unit was recorded (MEG: diamond; PEG: cross; AEG: circle; uncertain field: triangle) and marker size indicates whether cooling was determined to have a significant effect on firing rate in the two-step lasso GLM statistical tests (larger size indicates significance). B) The same plot as A, with axes limits expanded to show outlying units.

Next, we asked how multisensory response properties varied by cortical layer, and whether the effects of cooling were uniform across depths. Units were assigned to supragranular, granular and infragranular layers. While we found visual neurons at all cortical depths, they were most common, and least likely to be impacted by cooling, in the infragranular layers.

Our data demonstrate that a proportion of units’ visual response properties altered with cortical cooling. However, it is clear from Fig.4 that a sizeable proportion of auditory responses were also altered by cooling. Auditory units’ firing rates in response to sound could increase or decrease during cooling. While this could indicate that our cooling was not spatially restricted to visual cortex, there are a number of reasons why we do not believe this to be the case: firstly our temperature measurements across auditory cortex strongly indicated this was not the case. Secondly, our recordings in visual cortex demonstrated that at distances over a few 100 um away from the cooling loop visual firing was restored to normal (Atilgan et al., 2018). Finally, increased responses are not consistent with a general suppression of firing rate. To better understand the impact of cooling on auditory cortical neuron response properties, for each neuron, we calculated baseline firing rates during the full pre-cooling block and then calculated the z-scored firing rate in response to either auditory or visual stimuli both before and during cooling.

Stimulus specific changes in firing rate were visualized in a 2D space (Fig.4A). From the dense cluster of points around x < 0 and y < 0 we can see that many neurons do not change their firing rate, or show a mild suppression of firing in both stimulus conditions when visual cortex was cooled. Visual units are largely clustered within a triangle bounded by x = 0, falling below the y = 0 line and above x = y, indicating cooling more specifically impacts their firing rate during visual rather than auditory trials. This indicates a stimulus-evoked specificity to the impact of cooling, rather than what one might expect if effects were due to general firing rate suppression caused by excess cooling.

Similarly, auditory units cluster along y=0 with many falling below x = y, indicating that some auditory neurons show significant suppression (and others enhancement), of a similar magnitude to the visual units during cooling, and that this tends to be specific to firing on auditory trials rather than across both stimulus types. AV units show a more diverse pattern of effects; while the majority cluster along either axis, or within similar regions as the auditory and visual units rather than the x=y line, consistent with stimulus specific effects of cooling, a few lie close to the x = y line, indicating that their firing in both modalities is equally impacted by cooling.

We next considered the latencies of responses recorded in each field, in response to each modality of stimulus (Fig.5). As expected, auditory response latencies were shorter in the MEG than in the PEG and AEG, and visual response latencies were longer than auditory response latencies across all fields. We compared the response latencies of auditory/visual units that either were or were not influenced by a visual/auditory stimulus and saw that in MEG the visual neurons (some of which had very long latency responses occurring as late as 300 + ms) had longer latencies than audiovisual neurons, whereas in the PEG and MEG the opposite trend was observed. Across all three fields auditory units that were influenced by visual stimuli had slightly longer latencies than auditory (only) units.

**Figure 5.**
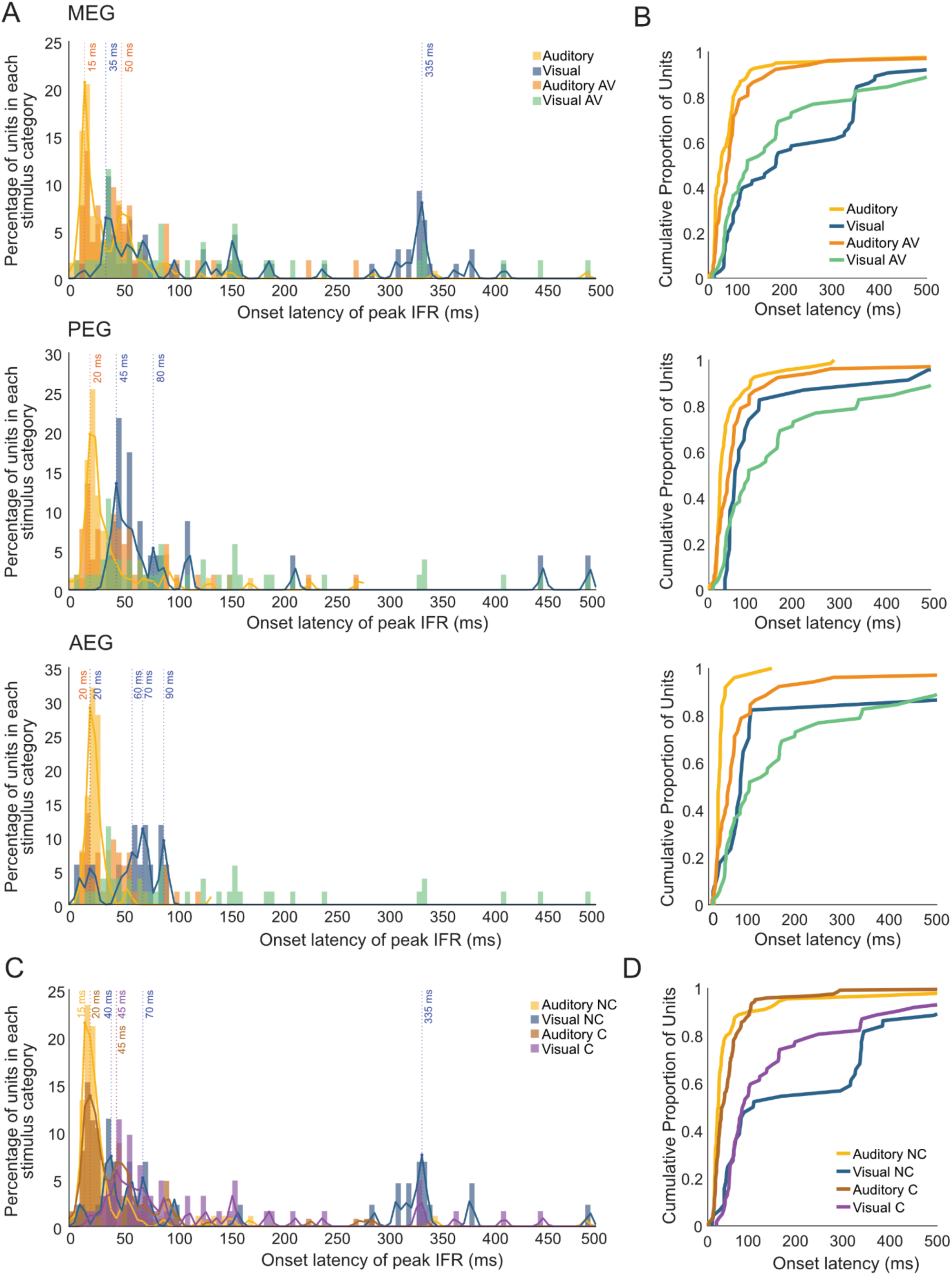
Latencies to peak instantaneous firing rate for auditory and visual responsive units. A) Histogram of percentages of units with onset latencies of peak instantaneous firing rate (IFR), calculated as latency to half-peak height in 5ms bins, separated by the recorded field into plots for MEG, PEG and AEG. Each plot shows four histograms, colour coded for units in four non-exclusive stimulus responsivity categories: auditory latencies for all auditory units, including A+V units (yellow), auditory latencies for A+V units (orange), visual latencies for all visual units, including A+V units (blue); visual latencies for only A+V units (green). Kernel density estimates (KDE) are plotted over the top of the auditory latencies for all units with an auditory response (yellow) and the visual latencies for all units with a visual response (blue). Peaks in the KDE are indicated in orange for the auditory latencies and blue for the visual latencies. B) Empirical cumulative distribution functions for the latencies plotted in A. C) Histogram of percentages of units with onset latencies of peak instantaneous firing rate (calculated as latency to half-peak height) in 5ms bins, pooled across all recorded fields. The plot shows four histograms, colour coded for units in four non-exclusive stimulus responsivity and cooling effect categories: auditory latencies for all auditory units not impacted by cooling (NC), including A+V units (yellow), visual latencies for all visual units not impacted by cooling, including A+V units (blue); auditory latencies for all auditory units impacted by cooling (C), including A+V units (brown), visual latencies for all visual units impacted by cooling, including A+V units (purple). Kernel density estimates (KDE) are plotted over the top for all four sets of latencies. Peaks in the KDE and their values are indicated in the corresponding colours. D) Empirical cumulative distribution functions for the latencies plotted in C.

Finally, we considered the response latencies of units that were, or were not, influenced by cooling (Fig.5C). Auditory units that were influenced by cooling had slightly longer latencies than auditory units that were uninfluenced by cooling. Visual units that were uninfluenced by cooling tended to have either shorter, or longer, latencies than those units that were affected by cooling, with the first peak of the KDE function for non-impacted units being around 40 ms as compared to 45 ms for the impacted units. The very late peak at 335 ms was only seen in visual units not impacted by cooling. The observation that units that were influenced or not by cooling have different latencies is potentially interesting, given that we would anticipate responses that originate sub-cortically may be either shorter (if they are direct) or much longer, if they form part of a cortical-subcortical-cortical loop.

## Discussion

We recorded neural responses to auditory, visual and audiovisual stimuli across auditory cortex while cooling areas PMLS and 21 in the visual cortex. We observed that roughly one third of visual responses were impacted by cooling, with the majority of responses being suppressed or eliminated. Our data are therefore consistent with PMLS/area 21 being one source of visual activity to auditory cortex.

In a handful of cases we observed that cooling unmasked a visual response that was previously absent, and which went away when visual cortical activity was restored. Such observations are hard to reconcile with a purely monosynaptic role of PMLS/area 21 and suggest instead that cooling PMLS/area 21 alters a third area which ultimately results in the unmasking of a usually inhibited or subthreshold visual response. Such circuit level effects should not be surprising, and unpicking whether this intermediary area is cortical or subcortical remains to be elucidated. While our data are consistent with a direct connection from PMLS/area 21 to auditory cortex providing visual responses to roughly half of the auditory cortical neurons that demonstrate visual sensitivity, to unequivocally demonstrate this would require pathway specific manipulation. For example, although silencing somatosensory cortex alters somatosensory responses in auditory cortex, the circuit basis for this is a cortex-midbrain-thalamus-cortex loop (Lohse et al., 2021). In the ferret, intersectional viral techniques would be required to accomplish pathway specific optogenetic manipulation. There are specific challenges involved with employing these more nuanced optogenetic techniques in the ferret, including infusing sufficient virus and generating the large area of illumination required in larger brained animals.

Contrary to our hypothesis, we saw that the effect of cooling was not restricted to the field that has the strongest input from PMLS/area 21. While AEG receives the strongest innervation from PMLS, PEG does receive direct innervation from PMLS, and area 21 in particular is interconnected with area 20, which projects heavily to the PEG (although note that when we made a small number of recordings ventral to the cooling loop – in area 20 – we did not observe an impact of cooling on these neurons). We have used the term ‘PMLS and area 21’ here as while our target – based on anatomical connectivity - was PMLS we could not cool within the sulcus without impacting auditory cortex directly, and without retinotopically mapping visual cortex we cannot be sure whether our cooling is restricted to the gyral surface of PMLS or encroaches on the anterior edge of area 21, which itself does not strongly project to auditory cortex, and lies within the lateral sulcus and suprasylvian gyrus. Given that previous work found many of the retrograde labelled cells from AEG lie deep within the sulcus it is therefore possible that our data underestimate the contribution of PMLS to visual responses in AEG, due to being unable to cool deep enough to impact the full population of cells projecting from PMLS to AEG. Finally, we note that the sample size of units we were able to track through the whole cooling cycle in AEG was rather small, imposing limitations on the strength of conclusions that we can draw with respect to AEG.

If PMLS provides only a proportion of the visual inputs to auditory cortex, where do the remainder originate from? Of particular note when considering this question are examples (such as Fig.2D) where one element of the visual response is lost during cooling, while others are maintained. We observed that loss during cooling was most likely to occur for longer latency response components raising the intriguing possibility that multiple visual inputs converge onto the same auditory cortical neurons. Other sources of (short latency) visual input include multisensory thalamus (such as suprageniculate nucleus and LP pulvinar). Within such a framework disinhibition via the thalamic reticular nucleus would also provide a potential explanation for the emergence of short latency visual responses during cooling.

We observed that audiovisual and visual responses were most common in the infragranular layers; this mirrors similar observations in the mouse (Morrill & Hasenstaub, 2018). Previous work in the ferret highlighted the supragranular layers as containing the most audiovisual responses (Atilgan et al., 2018). However, that study used very different stimuli (long luminance/amplitude modulated streams, as compared to the short 100 ms noise / light bursts used here), and didn’t look explicitly at visual responses as we have here, potentially providing an explanation for the difference.

The recordings reported here were made under anesthesia. It is possible, and indeed likely, that brain state, behavioural goals, experience and attention may all modulate the incidence of multisensory responses in auditory cortex (Garner & Keller, 2021). Nonetheless, studies that have explicitly compared anesthetised to passively listening awake animals have reported that similarities outweigh differences (Atilgan et al., 2018). One notable difference in awake animals was a higher proportion of audiovisual responses in the PEG, potentially offering an explanation for the counter-intuitive finding that primary auditory cortex contained a higher proportion of responses than PEG.

In conclusion, our study provides evidence that visual responses in ferret auditory cortex emerge from a variety of sources, and that one key area is a network that includes visual field PMLS/area 21.

## Acknowledgments

This work was supported in whole or in part by the Wellcome Trust through a Wellcome Trust/Royal Society Sir Henry Dale Fellowship Grant 098418/Z/12/A and a Career Development Award (227480/Z/23/Z to JKB, the BBSRC grant BB/H016813/1, a Royal Society Dorothy Hodgkin Fellowship and an ERC Consolidator award (771550, SOUNDSCENE).

We are grateful to Gareth Jones and Huriye Atilgan for assistance with data collection. We also wish to thank Stephen David and Jereme Wingert for sharing an early version of the visualisation toolbox we used as part of our layer analyses.

